# Metabolomics survey of uropathogenic bacteria in human urine

**DOI:** 10.1101/2024.10.07.617107

**Authors:** Carly C.Y. Chan, Ryan A. Groves, Thomas Rydzak, Ian A. Lewis

## Abstract

Urinary tract infections (UTIs) are one of the most prevalent infections in North America and are caused by a diverse range of bacterial species. Although uropathogenesis has been studied extensively in the context of macromolecular interactions, the degree to which metabolism may contribute to infection is unclear. Currently, most of what is known about the metabolic capacity of uropathogens has been derived from genomics, genetic knockout studies or transcriptomic analyses. However, there are currently very little empirical data on the metabolic activity of uropathogens when grown in urine. To address this gap, we conducted a systematic survey of the metabolic activities of eight of the most common uropathogenic bacterial species that collectively represent 99% of uncomplicated UTIs. Liquid chromatography-mass spectrometry (LC-MS) analyses of human urine cultures revealed that uropathogens have four distinct metabolic clades. We generalized these clades as serine consumers (*Escherichia coli, Klebsiella pneumoniae* and *Proteus mirabilis*), glutamine consumers (*Pseudomonas aeruginosa*), amino acid abstainers (*Enterococcus faecalis* and *Streptococcus agalactiae*), and amino acid minimalists (*Staphylococcus aureus* and *Staphylococcus saprophyticus*). These metabolic classifications can be further subdivided on a species-to-species level. This survey provides a framework to understanding the metabolic activity of the diverse range of uropathogens and how these species use divergent metabolic strategies to occupy the same niche.

## 1 Introduction

Urinary tract infections (UTIs) are one of the most common infections, and are responsible for 8 million healthcare visits each year in the United States alone (Schappert and Rechtsteiner, 2011). UTIs can be caused by a diverse range of microorganisms, but in an outpatient setting, eight bacterial species are responsible for 99% of infections (Flores-Mireles et al., 2015; Wagenlehner et al., 2020). This relatively small number of species is somewhat surprising given the significant diversity of species identified in human feces, the presumed source of uropathogens (Tannock, 1999; Segata et al., 2012). This suggests that selective forces are at play, and one contributing factor may be nutritional selection.

Most of what we know about UTIs comes from *Escherichia coli*, which is frequently used as the model pathogen in studies since it is responsible for around 75% of uncomplicated UTIs (Flores-Mireles et al., 2015; Wagenlehner et al., 2020). Information about uropathogenic *E. coli* metabolism comes from genomic mapping, genetic knockouts, or transcriptional studies (Chan and Lewis, 2022). The consensus in the literature is that uropathogenic *E. coli* must take up a wide range of urinary amino acids to fuel central carbon metabolism (Alteri et al., 2009; Chan and Lewis, 2022). In particular, serine catabolism is essential for *E. coli* growth in urinary environments (Hull and Hull, 1997; Roesch et al., 2003; Anfora and Welch, 2006; Anfora et al., 2007). Uropathogenic *E. coli* has also been shown to salvage various nucleotides, presumably to support gene replication (Russo et al., 1996; Vejborg et al., 2012; Shaffer et al., 2017; Andersen-Civil et al., 2018; Ma et al., 2018). In addition, *E. coli* also relays on urinary ethanolamine as a nitrogen source (Sintsova et al., 2018; Dadswell et al., 2019).

Although the metabolic requirements for uropathogenic *E. coli* are now emerging, this is only one of the many uropathogenic species, and thus, represents an incomplete picture of the metabolic selective strategies that could contribute to infection. Other uropathogenic species have yet to be metabolically characterized, despite collectively accounting for the remaining 25% of infections (Flores-Mireles et al., 2015; Wagenlehner et al., 2020). These species include *Klebsiella pneumoniae, Staphylococcus saprophyticus, Enterococcus faecalis, Streptococcus agalactiae* (group B streptococcus), *Proteus mirabilis, Pseudomonas aeruginosa, Staphylococcus aureus*, and *Candida spp*. (Flores-Mireles et al., 2015; Wagenlehner et al., 2020). Of these rarer uropathogens, infections caused by *P. mirabilis* have been attracting an increasing degree of attention because of its ability to produce crystalline biofilms which leads to complicated catheter-associated UTIs (Norsworthy and Pearson, 2017; Gmiter and Kaca, 2022). Similar to *E. coli, P. mirabilis* was also found to preferentially catabolize serine in human urine (Brauer et al., 2019, 2022). However, the metabolic needs of all uropathogens outside of *E. coli* is poorly understood. Moreover, there have not been any in-depth investigations of the metabolic distinctions between different uropathogenic species.

Recent advances of high-resolution liquid chromatography-mass spectrometry (LC-MS) have radically expanded the complement of metabolic activities that can be tracked in routine studies. Moreover, our group has recently developed a specialized metabolic boundary flux analysis strategy used to characterize the metabolic phenotypes of microbe based on the rates of nutrient uptake and waste product excretion (Lewis, 2024). These metabolic profiles provide a convenient mechanism for making interspecific comparisons of metabolic activities and are useful for identifying metabolic phenotypes that distinguish closely related species (Rydzak et al., 2022). Herein, we harnessed this boundary flux analysis strategy to conduct a systematic metabolomics survey of the eight most common uropathogenic species to provide a systematic review of the nutritional strategies used by these species when grown in human urine.

## 2 Materials and methods

### 2.1 Human urine collection and bacterial strains

Human urine was collected and pooled from five adult donors (three females, two males), following institutional ethics board approval (REB19-0442). Immediately after collection, the pooled urine stock was filtered sterilized, aliquoted, and stored at -20 °C. In preparation for experiments, aliquots of the pooled urine were thawed at room temperature and centrifuged (4,200 × g, 15 min). This pooled stock was used for all experiments presented in this study.

Bacterial species most frequently responsible for uncomplicated UTIs—each contributing to at least 1% of infections in outpatient setting—were selected for our metabolomics survey. A total of eight species were evaluated including four Gram-negative species (*E. coli, K. pneumoniae, P. aeruginosa*, and *P. mirabilis*) and four Gram-positive species (*E. faecalis, S. agalactiae, S. aureus*, and *S. saprophyticus*). For each species, six diverse strains were selected from different isolation origins, including laboratory controls and clinical isolates from UTIs and other infections (Table S1).

### 2.2 *In vitro* bacteria growth in pooled human urine

A total of 48 bacterial strains were grown *in vitro* in pooled, sterile-filtered human urine. All bacterial strains were first cultured overnight in Mueller Hinton (MH) medium (BD Difco, Mississauga, ON, Canada) at 37 °C with shaking (180 rpm). Overnight cultures were centrifuged (4,200 × g, 15 min) and washed twice with phosphate buffered saline (PBS) to remove contaminating metabolites. Strains were then inoculated at an OD_600_ (optical density at 600 nm) of 0.1 in pooled human urine (Mutiskan™ GO Microplate Spectrophotometer, Thermo Scientific), producing 200 µL bacterial cultures on a 96-well plate. Thus, all urine cultures were normalized to approximately 1 × 10^5^ CFU/mL at the start of the experiment. The urine cultures were incubated at 37 °C with 5% CO_2_ and 21% O_2_ (Heracell™ VIOS 250i Tri-Gas Incubator, Thermo Scientific) for four hours and the OD_600_ of the cultures were monitored at 30-minute intervals (Mutiskan™ GO Microplate Spectrophotometer, Thermo Scientific).

Extracellular samples of the cultures were taken at the start and end of the growth course, and then diluted in HPLC (high-performance liquid chromatography)-grade methanol at a 1:1 ratio and frozen at -80 °C. Additionally, at the end of the growth course, the cultures were plated on MH agar plates (except for *S. agalactiae* cultures that were plated on Columbia blood agar instead) at 10^5^ and 10^6^ dilutions. The agar plates were incubated overnight at 37 °C, and then the resultant colonies were counted.

### 2.3 Liquid chromatography-mass spectrometry analysis

LC-MS analysis was conducted at the Calgary Metabolomics Research Facility. In preparation for metabolomics analysis, extracellular samples collected from urine cultures were thawed at room temperature, centrifuged (4,200 × g, 10 min), and diluted ten-fold with 50% HPLC-grade methanol before undergoing LC-MS analysis. Our LC-MS methods has been described in detail elsewhere (Groves et al., 2022; Rydzak et al., 2022). Briefly, metabolites in the samples were resolved with a Syncronis™ Hydrophilic Interaction Liquid Chromatography (HILIC) Column (2.1 mm × 100 mm × 1.7 µm, Thermo Scientific) on a Vanquish™ Ultra-High-Performance Liquid Chromatography (UHPLC) platform (Thermo Scientific) using a 15-minute two-solvent gradient method (20 mM ammonium formate at pH 3.0 in HPLC-grade water and HPLC-grade acetonitrile with 0.1% formic acid). Mass spectrometry data were acquired in full scan in both positive and negative modes on a Q Exactive™ HF Hybrid Quadrupole-Orbitrap™ Mass Spectrometer (Thermo Scientific). All acquired LC-MS data were analyzed with El-MAVEN v0.12.0 software (Agrawal et al., 2019).

Metabolites were assigned using an in-house library of chemical standards on the basis of exact mass and chromatographic retention time. Each assignment was verified using a commercial standard from Sigma-Aldrich. Chemical standards were also used to prepare a concentration calibration reference mixture for absolute quantification, and metabolite concentrations were computed following established methods (Ponce et al., 2024). Metabolic activity exhibited by each species was identified based on the consumption or production of metabolites. These were defined as threshold differences relative to uninoculated urine controls. Significantly consumed or secreted metabolites were identified using Welch’s two-sample t-test followed by Bonferroni correction (α=0.00625).

### 2.4 Characterization of a cohort of clinical polymicrobial urine samples

To establish the frequency of which certain species co-segregated in polymicrobial UTIs, we acquired the species information of a cohort of patient urine samples sent to Alberta Precision Laboratories over the course of a week. The demographics of UTIs from the Calgary health region has been previously described (Laupland et al., 2007; Gregson et al., 2021). Out of 1,462 urine samples identified as growth-positive, only 81 samples contained two causative species, and the species information was collected from this subset. To determine if the observed species distribution was expected by chance, the top three pairs of species underwent individual chi-square goodness-of-fit tests.

## 3 Results

### 3.1 Metabolic composition of human urine

The chemical composition of human urine can vary considerably between individuals and over time (Rasmussen et al., 2011; Bouatra et al., 2013; Virgiliou et al., 2021). Thus, to minimize variability, urine samples were pooled into one stock and this stock was used for all experiments. As expected, metabolomics analysis of our pooled urine stock revealed that it contained high concentrations of amino acids and relatively low concentrations of carbohydrates and nucleic acids (Table S2). The concentrations of individual metabolites in our pooled urine stock were within the normal ranges reported in the literature, with a few exceptions including isoleucine, leucine, cystine, 4-hydroxyproline, uracil, and succinate, which were approximately 10 µM more or less abundant than published normal ranges (Bouatra et al., 2013).

### 3.2 Metabolomics analysis and identification of metabolic clades

For each species, six distinct isolates were seeded at 1 × 10^5^ CFU/mL in pooled human urine and grown for four hours, reaching 3.0 × 10^6^ to 3.6 × 10^8^ CFU/mL. The metabolic profiles observed in the cultures over time were captured by LC-MS using our previously established strategy (Rydzak et al., 2022; Lewis, 2024) (Figure 1). The metabolite abundances in the cultures were compared to those in the urine controls to determine changes in metabolite concentration and identify metabolites that were consumed or produced (Figure S1 and Table S3). As expected, many of the metabolic phenotypes that we observed were consistent with those reported in the literature. For example, *E. coli* is well-known for utilizing glucose as its preferred carbon source and secreting succinate (Holms, 1996; Thakker et al., 2012), a phenotype which we also observed when it was grown in urine (Table S3). Similarly, tyramine production is well-characterized in *E. faecalis* (Connil et al., 2002; Perez et al., 2015), and we also observe its production in *E. faecalis* urine cultures (Table S3).

**Figure 1.**
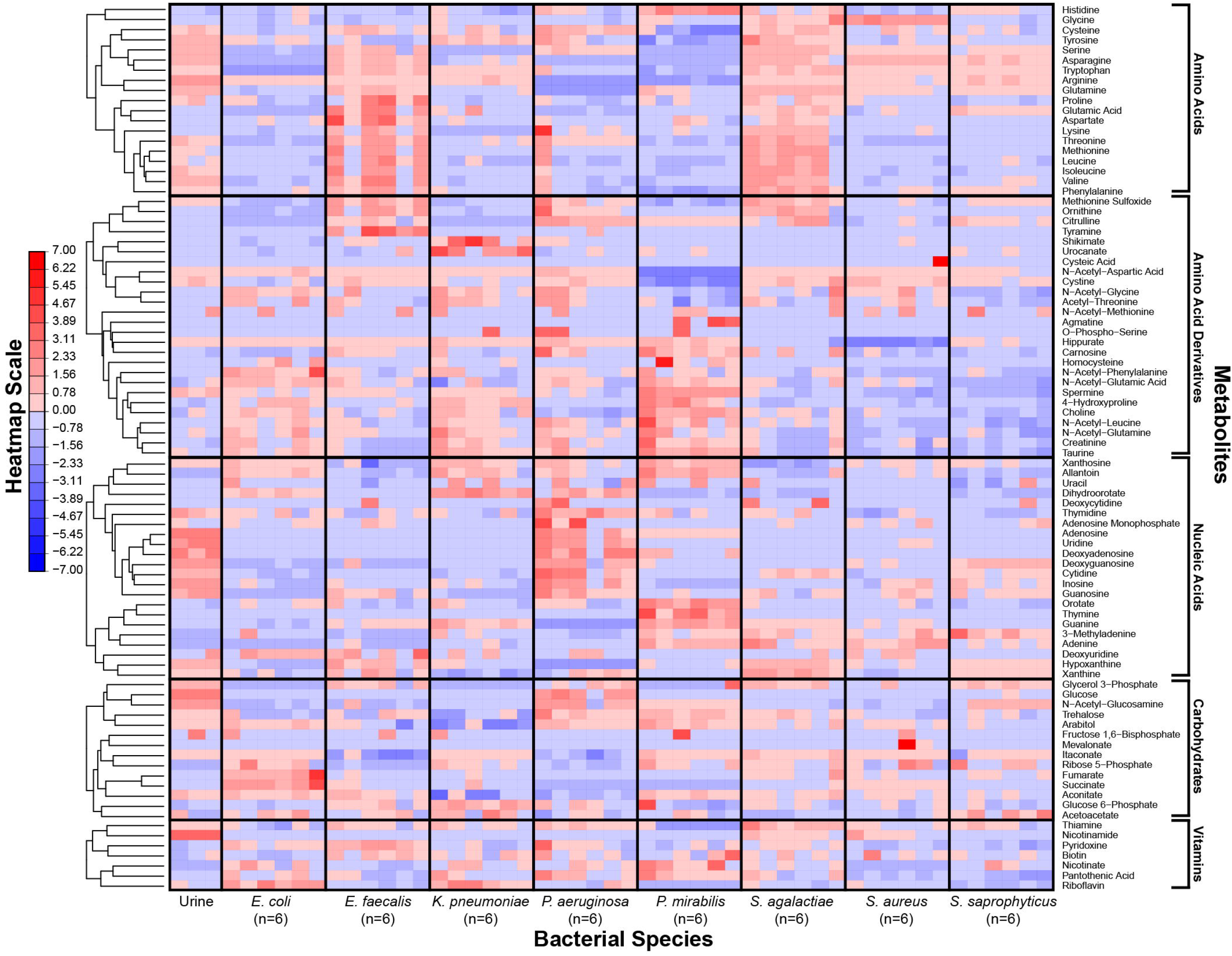
Metabolic phenotypes of uropathogens when grown in human urine. Metabolite concentrations in *in vitro* cultures were quantified by LC-MS after four-hour incubation in pooled human urine. For each species, six distinct isolates were used to assess phenotypic diversity. Differences in metabolite concentrations across species are depicted as z-scores. Metabolite classes are listed (right) and metabolic phenotypes are clustered according to hierarchical clustering (left). Red indicates higher metabolite concentration and blue indicates lower metabolite concentration.

Our metabolomics approach enabled us to capture a more comprehensive transect of metabolites across our panel of microorganisms. As reported previously (Rydzak et al., 2022), bacterial strains of the same species showed consistent metabolic phenotypes (Figure 1). However, each species exhibited a unique metabolic profile of consumed versus secreted metabolites (Figure S1). Multivariate analyses of the metabolic profiles observed in these uropathogens showed that although individual species-related clusters occurred, broader phenotypes were also observed across groups of species (Figure 2). As expected, the most significant metabolic differences were between Gram-negative species (*E. coli, K. pneumoniae*, and *P. mirabilis*) and Gram-positive species (*E. faecalis, S. agalactiae, S. aureus*, and *S. saprophyticus*) with *P. aeruginosa* being a notable exception that occupies a unique metabolic space distinct from all other uropathogens (Figure 2A). Based on similarities and differences in metabolic profiles, uropathogens were generalized into four metabolic clades: 1) serine consumers, 2) glutamine consumers, 3) amino acid abstainers, and 4) amino acid minimalists (Figure 2B). Each of these labels are proxies to describe a complex suite of metabolic phenotypes that generally clusters these uropathogenic species.

**Figure 2.**
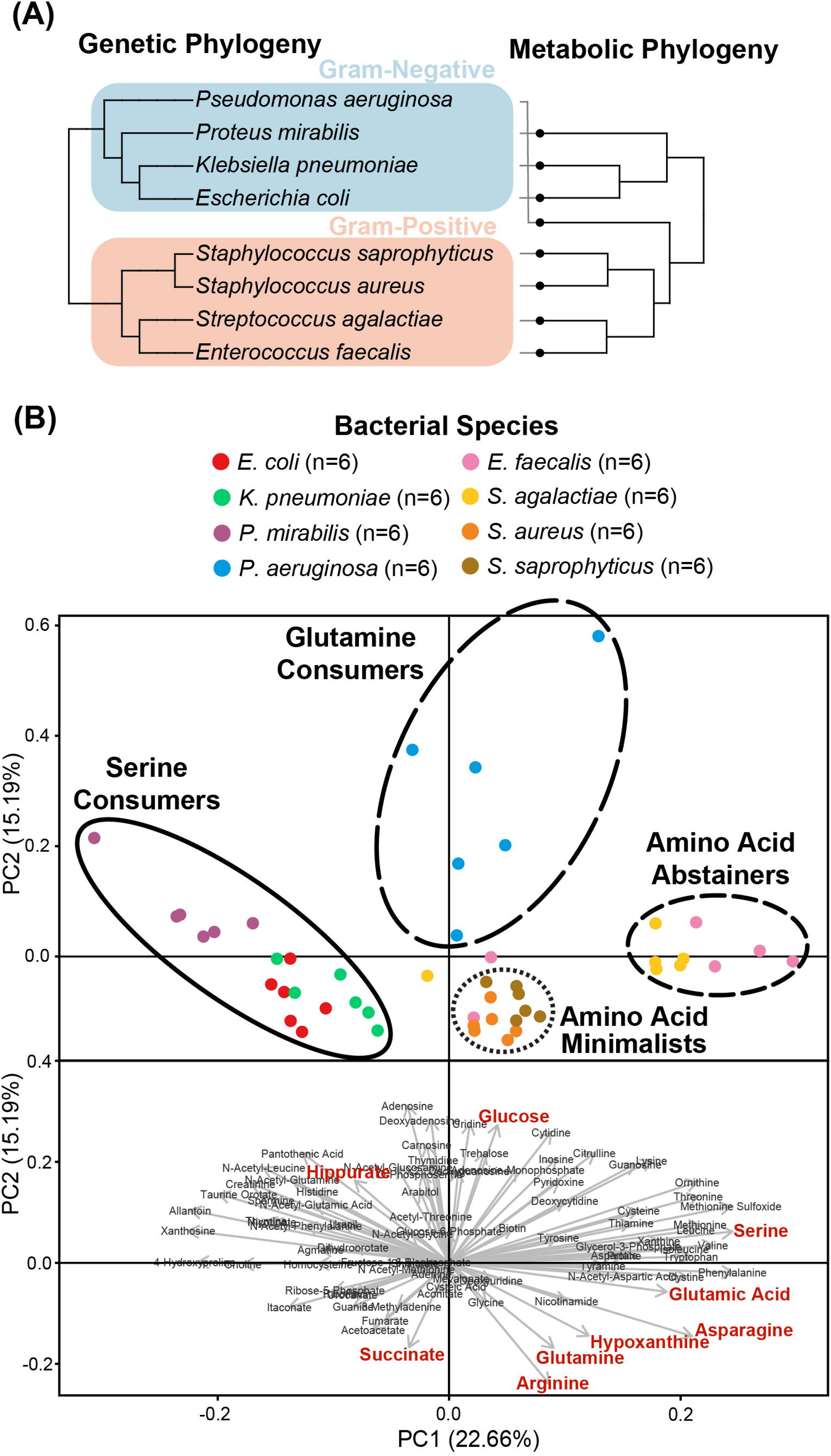
Clustering of the metabolic profiles of uropathogens. **(A)** Clustering of uropathogens according to their genomic and metabolic phylogenies. Genomic phylogenies were derived from phyloT v2 (https://phylot.biobyte.de/) and were compared to the metabolic classifications derived from hierarchical clustering. **(B)** Clusters of metabolic phenotypes from principal component analysis (PCA). Score plot distributions of eight species were shown with the metabolic clades illustrated by circles. PCA-biplot distribution of important metabolites that are contributing to the segregation of the scores. Red metabolites denote phenotypes that were identified as significant after Bonferroni correction in univariate analyses.

The cluster of “serine consumers” included *E. coli, K. pneumoniae* and *P. mirabilis*. All of these Gram-negative *Enterobacterales* species consumed between 90-98% of urinary serine (Figure 3A). *P. aeruginosa* is an outlier with regards to the Gram-negative group as it failed to consume serine (Figure 3A). *P. aeruginosa* was also the only species not to consume glucose, instead consuming higher concentrations glutamine (95%) and hypoxanthine (97%) (Figure 3A). Due to its distinct profile, *P. aeruginosa* was categorized into a metabolic clade of its own as a “glutamine consumer”, despite displaying a few shared phenotypes with *P. mirabilis*, such as high arginine and succinate consumption (Figure S2).

**Figure 3.**
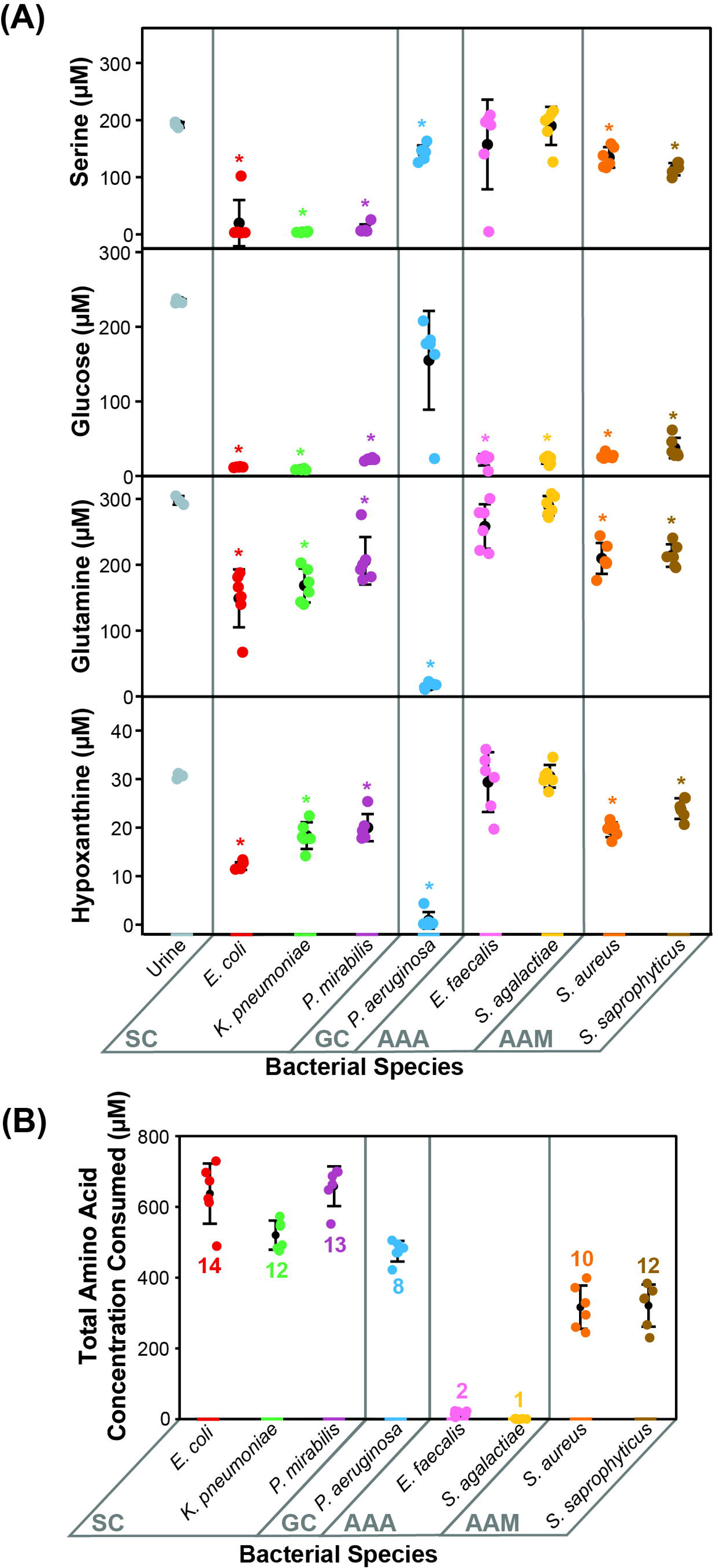
Prominent metabolic features that defined each metabolic clade. Metabolic phenotypes distinctive to **(A)** Gram-negative and **(B)** Gram-positive metabolic clades are displayed, including serine, glucose, glutamine, hypoxanthine, and the total amino acid concentration consumed along with number of different amino acids consumed. Statistical significance is denoted by an asterisk (^*^, two-sample t-test with Bonferroni correction α=0.00625). Error bars represent one standard deviation. Abbreviations: SC, serine consumers; GC, glutamine consumers; AAA, amino acids abstainers; AAM, amino acids minimalists.

The clusters of “amino acid abstainers” and “amino acid minimalists” were exclusively made up of Gram-positive organisms that consumed very few amino acids compared to Gram-negative species (Figure 3B). Of these, the amino acid abstainers (*E. faecalis* and *S. agalactiae*) only consumed between 1-2 amino acids out of the 19 amino acids detected, while other species across our cohort consumed between 8-14 (Figure 3B). Whereas, the amino acid minimalists (*S. saprophyticus* and *S. aureus*) consumed a broader range of amino acids than amino acid abstainers but at a lower quantity than the Gram-negative bacteria (Figure 3B).

Most of the bacterial species evaluated consumed 319.50-664.10 µM of the total amino acids present (Table S3). The exceptions were the amino acid abstainers (*E. faecalis* and *S. agalactiae*), which only consumed 22.04 µM and 6.24 µM of amino acids, respectively (Table S3). Nearly all observed amino acids (16/19 amino acids) were consumed by at least one bacterial species (Table S3). Urinary arginine, asparagine, glutamine, isoleucine, methionine, serine, threonine, tryptophan, tyrosine, and valine were consumed by over half of all species assessed (≥4/8 species) (Table S3). Neither glycine nor histidine were consumed by any species, despite being the top two most abundant amino acids in urine (Table S3). In contrast, the next two most abundant amino acids, serine and glutamine, which were consumed by six out of eight species (Table S3). As noted previously, *K. pneumoniae, P. mirabilis*, and *E. coli* consumed higher quantities of serine (90-98%), while *P. aeruginosa* consumed higher quantities of glutamine (95%) relative to other species (Table S3).

As expected, urinary glucose was consumed by seven out of the eight species, each consuming between 84-96% (Table S3). *P. aeruginosa* was the only exception, which did not significantly consume glucose (Figure 3A). Another prevalent carbon source in urine was succinate, and between 86-99% was consumed by *P. aeruginosa* and *P. mirabilis* (Figure S2 and Table S3). Interestingly, succinate was also produced by *E. coli* (1003.44 µM), *S. saprophyticus* (47.74 µM) and *S. aureus* (184.60 µM) (Figure S2 and Table S3).

Nearly all available urinary nucleosides (9/11 nucleosides) were taken up by most species (≥5/8 species), including adenosine, cytidine, deoxyadenosine, deoxyguanosine, guanosine, inosine, thymidine, and uridine (Table S3). In contrast, nucleobases were rarely consumed (2/6 nucleobases), except for hypoxanthine and xanthine (Table S3). A few species secreted nucleic acids, and the most pronounced example of this was the unique thymine production phenotype observed in *P. mirabilis* (203.62 µM) (Figure S2 and Table S3). This metabolic phenotype appeared to be specific to *P. mirabilis* urine culture, as it was not observed when it was grown in other growth media such as MH broth (Figures S3).

### 3.3 Prevalence of species in polymicrobial UTIs

A significant fraction of UTIs are polymicrobial (Gaston et al., 2021). Given the distinct metabolic clades observed in this study, we speculated that polymicrobial infections may be more prevalent in species with complimentary metabolic niches. To test this hypothesis, we collected the species breakdown of all polymicrobial infections observed in the Calgary health zone over a one-week period, and in this cohort, 5.5% were identified as polymicrobial (81/1,462). The top three most frequent pairings in this cohort were *E. coli* and *E. faecalis* (32.1%), *E. coli* and *K. pneumoniae* (8.6%), and *E. coli* and *Streptococcus viridans* (8.6%) (Table S4). To better understand if the co-segregation of these species followed the expected distribution based on their individual prevalence, we conducted chi-square goodness-of-fit tests on the top three pairs (Table S4). The *E. coli* and *E. faecalis* pair (p=0.0026) and the *E. coli* and *S. viridans* pair (p=0.014) occurred significantly more frequently than expected, while the prevalence of the *E. coli* and *K. pneumoniae* pair (p=0.48) followed the distribution expected by chance (Table S4).

## 4 Discussion

The primary goal of this study was to provide empirical evidence for the metabolic preferences of common uropathogens when grown in human urine. To the best of our knowledge, this is the first study that provides the foundational metabolic data needed to understand the diverse nutritional strategies used by these uropathogens. Our metabolomics data showed that the eight most common uropathogens followed four nutritional strategies which we characterized as serine consumers, glutamine consumers, amino acid abstainers, and amino acid minimalists (Figure 2B). These categorical designations are not exhaustive lists of the metabolic differences between the clades but serves as a convenient descriptors of the most distinctive metabolic features in each group (Figure 3). The nutritional strategies generally unique to each metabolic clade are illustrated in Figure 4.

**Figure 4.**
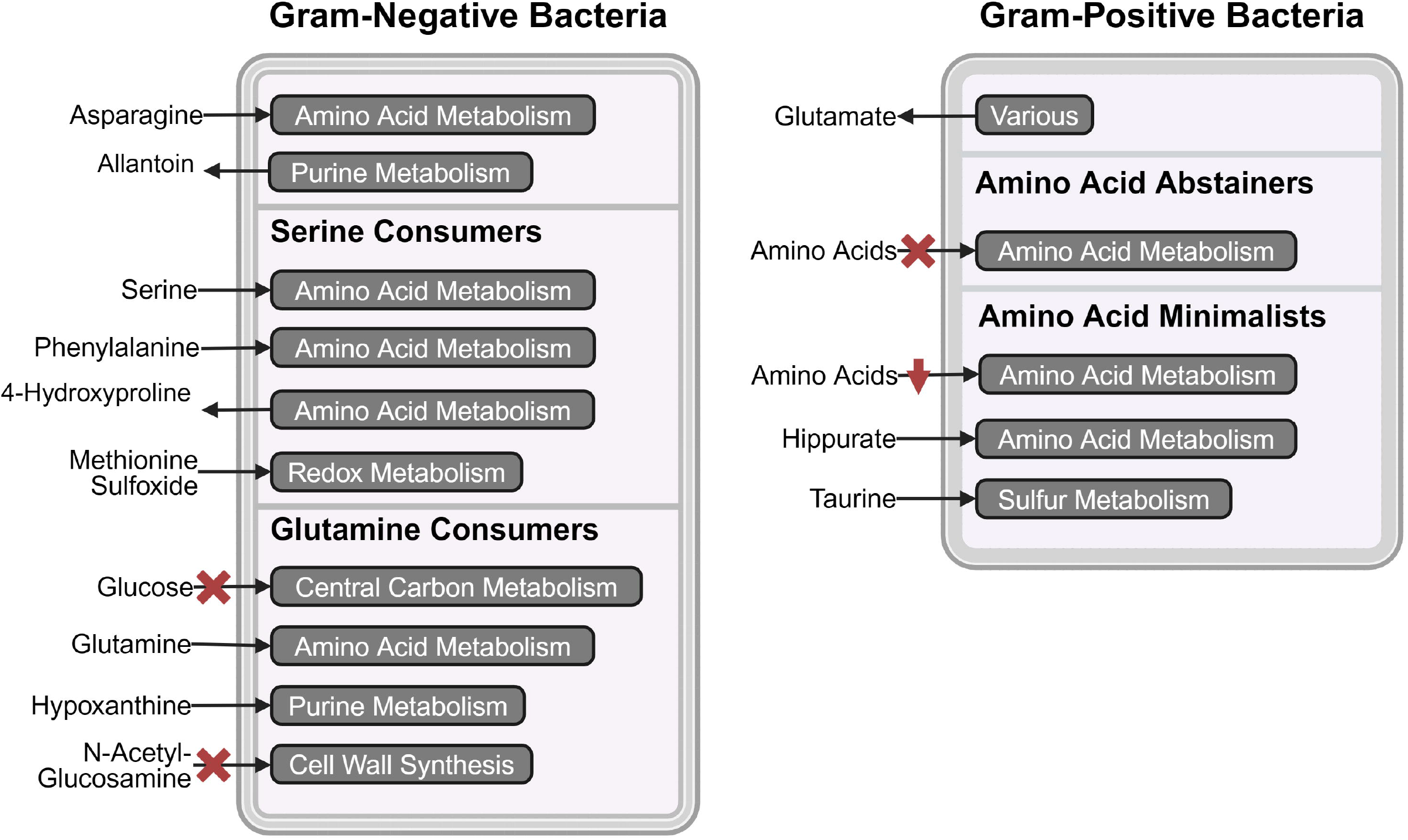
Schematic representation of metabolic pathway activities that distinguish the four metabolic clades. The most prominent consumed and secreted metabolites, and their respective metabolic pathways are shown for serine consumers, glutamine consumers, amino acid abstainers, and amino acid minimalists. The red crosses indicate the absence of the metabolic phenotype and red downward arrows indicate lower activity. Figure was made with BioRender.com.

As reported in similar studies (Rydzak et al., 2022), our metabolomics analysis showed that the metabolic features within strains of the same species were minimal relative to interspecific differences (Figure 1). The metabolic distinctions between species tended to co-segregate according to phylogeny (Figure 2A). The most dramatic metabolic distinction observed in this dataset were those that separated Gram-negative (*E. coli, K. pneumoniae, P. mirabilis*, and *P. aeruginosa*) and Gram-positive species (*E. faecalis, S. agalactiae, S. aureus*, and *S. saprophyticus*) (Figure 2B). Gram-negative species are evidently well-adapted to using a wide range of urinary amino acids, while the Gram-positive species appear to use little to no amino acid catabolism to support their growth (Figure 3B).

Divergence in nutritional strategies may provide a selective advantage for uropathogens, especially when competing against fast-growing organisms such as *E. coli*. This may be relevant in the context of polymicrobial infections, wherein two or more species co-exist in the urinary tract (Azevedo et al., 2017; De Vos et al., 2017; Gaston et al., 2021). Our analysis of the species breakdown of polymicrobial infections in the Calgary health zone appears to support this hypothesis. We observed significant co-segregation of *E. coli* and *E. faecalis* (p=0.0026, chi-square goodness-of-fit test) (Table S4) and this pairing was the most prevalent in other documented cohorts as well (Cardone et al., 2018; Folaranmi et al., 2022). This significant co-segregation may occur because *E. coli* and *E. faecalis* occupy compliment metabolic niches with *E. coli* as a serine consumer and *E. faecalis* as an amino acid abstainer. *E. coli* depends heavily on amino acid catabolism, consuming 14 out of 19 amino acids totaling to 643.43 µM, whereas *E. faecalis* does not, consuming only two amino acids totaling to 22.04 µM (Figure 3B and Table S3). This metabolic distinction may allow *E. faecalis* to establish infections in the urinary tract without competing in the same metabolic niche as *E. coli*.

One interesting application of our dataset is as a tool to help refine artificial urine. Artificial urine is used routinely to create a more physiological environment for *in vitro* studies of urinary tract diseases (Grases et al., 1996; Jacobs et al., 2001; Jones et al., 2007; Ma et al., 2018; Juarez et al., 2020). However, we note that the metabolic compositions in current artificial urine formulations are neither defined nor reflective of the metabolite abundances observed in human urine (Brooks and Keevil, 1997; Chutipongtanate and Thongboonkerd, 2010; Ipe et al., 2016; Sarigul et al., 2019). This is problematic because many of the metabolites we observed as preferred nutrients (outlined in Table S2) are absent or not chemically defined in established formulas. Therefore, supplementing artificial urine with these key metabolite can potentially help bring these formulas closer to physiological relevance.

In summary, we conducted a systematic metabolomics survey of the eight most common uropathogens and identified four distinct metabolic clades that differentiate uropathogens. We observed significant differences in nutritional styles that may help explain the co-segregation of species in polymicrobial infections and pave the way for creating a more precisely defined artificial urine media. Overall, this dataset provides a foundation for future metabolomics analyses of uropathogens and clearly demonstrated that the nutritional strategies of uropathogens should be considered in future UTI studies.

## 6 Author Contributions

CCYC designed and performed the experiments, interpreted the mass spectrometry data, and wrote the manuscript. RAG and TR developed microbial growth protocols and mass spectrometry methods. IAL provided advice and feedback on the research project and manuscript. All authors contributed to the article and approved the submitted version.

## 8 Acknowledgments

Special thanks to Dr. Daniel B. Gregson for providing us with data on the prevalence of different species in polymicrobial UTIs. We also thank Dr. Stephanie Bishop for providing the chemical standard mixes for quantitative LC-MS analysis, and Keir Pittman for critical review and editorial assistance in preparing this manuscript.

## References

Agrawal, S., Kumar, S., Sehgal, R., George, S., Gupta, R., Poddar, S., et al. (2019). “El-MAVEN: A Fast, Robust, and User-Friendly Mass Spectrometry Data Processing Engine for Metabolomics,” in High-Throughput Metabolomics: Methods and Protocols, ed. A. D’Alessandro (Methods in Molecular Biology), 301–321. doi: 10.1016/B978-1-4160-3106-2.00019-2

Alteri, C. J., Smith, S. N., and Mobley, H. L. T. (2009). Fitness of Escherichia coli during urinary tract infection requires gluconeogenesis and the TCA cycle. PLoS Pathog. 5, e1000448. doi: 10.1371/journal.ppat.1000448

Andersen-Civil, A. I. S., Ahmed, S., Guerra, P. R., Andersen, T. E., Hounmanou, Y. M. G., Olsen, J. E., et al. (2018). The impact of inactivation of the purine biosynthesis genes, purN and purT, on growth and virulence in uropathogenic E. coli. Mol. Biol. Rep. 45, 2707–2716. doi: 10.1007/s11033-018-4441-z

Anfora, A. T., Haugen, B. J., Roesch, P., Redford, P., and Welch, R. A. (2007). Roles of serine accumulation and catabolism in the colonization of the murine urinary tract by Escherichia coli CFT073. Infect. Immun. 75, 5298–5304. doi: 10.1128/IAI.00652-07

Anfora, A. T., and Welch, R. A. (2006). DsdX is the second D-serine transporter in uropathogenic Escherichia coli clinical isolate CFT073. J. Bacteriol. 188, 6622–6628. doi: 10.1128/JB.00634-06

Azevedo, A. S., Almeida, C., Melo, L. F., and Azevedo, N. F. (2017). Impact of polymicrobial biofilms in catheter-associated urinary tract infections. Crit. Rev. Microbiol. 43, 423–439. doi: 10.1080/1040841X.2016.1240656

Bouatra, S., Aziat, F., Mandal, R., Guo, A. C., Wilson, M. R., Knox, C., et al. (2013). The human urine metabolome. PLoS One 8, e73076. doi: 10.1371/journal.pone.0073076

Brauer, A. L., Learman, B. S., Taddei, S. M., Deka, N., Hunt, B. C., and Armbruster, C. E. (2022). Preferential catabolism of L-vs D-serine by Proteus mirabilis contributes to pathogenesis and catheter-associated urinary tract infection. Mol. Microbiol. 118, 125–144. doi: 10.1111/mmi.14968

Brauer, A. L., White, A. N., Learman, B. S., Johnson, A. O., and Armbruster, C. E. (2019). D-Serine degradation by Proteus mirabilis contributes to fitness during single-species and polymicrobial catheter-associated urinary tract infection. mSphere 4, e00020–19. doi: 10.1128/mSphere.00020-19

Brooks, T., and Keevil, C. W. (1997). A simple artificial urine for the growth of urinary pathogens. Lett. Appl. Microbiol. 24, 203–206.

Cardone, S., Petruzziello, C., Migneco, A., Fiori, B., Spanu, T., D’Inzeo, T., et al. (2018). Age-related trends in adults with urinary tract infections presenting to the emergency department: A 5-year experience. Rev. Recent Clin. Trials 14, 147–156. doi: 10.2174/1574887114666181226161338

Chan, C. C. Y., and Lewis, I. A. (2022). Role of metabolism in uropathogenic Escherichia coli. Trends Microbiol. 30, 1174–1204. doi: 10.1016/j.tim.2022.06.003

Chutipongtanate, S., and Thongboonkerd, V. (2010). Systematic comparisons of artificial urine formulas for in vitro cellular study. Anal. Biochem. 402, 110–112. doi: 10.1016/j.ab.2010.03.031

Connil, N., Le Breton, Y., Dousset, X., Auffray, Y., Rincé, A., and Prévost, H. (2002). Identification of the Enterococcus faecalis tyrosine decarboxylase operon involved in tyramine production. Appl. Environ. Microbiol. 68, 3537–3544. doi: 10.1128/AEM.68.7.3537-3544.2002

Dadswell, K., Creagh, S., McCullagh, E., Liang, M., Brown, I. R., Warren, M. J., et al. (2019). Bacterial microcompartment-mediated ethanolamine metabolism in Escherichia coli urinary tract infection. Infect. Immun. 87, e00211–19. doi: 10.1128/IAI.00211-19

De Vos, M. G. J., Zagorski, M., McNally, A., and Bollenbach, T. (2017). Interaction networks, ecological stability, and collective antibiotic tolerance in polymicrobial infections. Proc. Natl. Acad. Sci. U. S. A. 114, 10666–10671. doi: 10.1073/pnas.1713372114

Flores-Mireles, A. L., Walker, J. N., Caparon, M., and Hultgren, S. J. (2015). Urinary tract infections: Epidemiology, mechanisms of infection and treatment options. Nat. Rev. Microbiol. 13, 269–284. doi: 10.1038/nrmicro3432

Folaranmi, T., Harley, C., Jolly, J., and Kirby, A. (2022). Clinical and microbiological investigation into mixed growth urine cultures. J. Med. Microbiol. 71, 001544. doi: 10.1099/jmm.0.001544

Gaston, J. R., Johnson, A. O., Bair, K. L., White, A. N., and Armbruster, C. E. (2021). Polymicrobial interactions in the urinary tract: Is the enemy of my enemy my friend? Infect. Immun. 89, e00652–20. doi: 10.1128/IAI.00652-20

Gmiter, D., and Kaca, W. (2022). Into the understanding the multicellular lifestyle of Proteus mirabilis on solid surfaces. Front. Cell. Infect. Microbiol. 12, 864305. doi: 10.3389/fcimb.2022.864305

Grases, F., Söhnel, O., Vilacampa, A. I., and March, J. G. (1996). Phosphates precipitating from artificial urine and fine structure of phosphate renal calculi. Clin. Chim. Acta 244, 45–67. doi: 10.1016/0009-8981(95)06179-7

Gregson, D. B., Wildman, S. D., Chan, C. C. Y., Bihan, D. G., Groves, R. A., Aburashed, R., et al. (2021). Metabolomics strategy for diagnosing urinary tract infections. medRxiv [Preprint]. Available at: 10.1101/2021.04.07.21255028 (Accessed October 1, 2024).

Groves, R. A., Mapar, M., Aburashed, R., Ponce, L. F., Bishop, S. L., Rydzak, T., et al. (2022). Methods for quantifying the metabolic boundary fluxes of cell cultures in large cohorts by high-resolution hydrophilic liquid chromatography mass spectrometry. Anal. Chem. 94, 8874–8882. doi: 10.1021/acs.analchem.2c00078

Holms, H. (1996). Flux analysis and control of the central metabolic pathways in Escherichia coli. FEMS Microbiol. Rev. 19, 85–116. doi: 10.1016/S0168-6445(96)00026-5

Hull, R. A., and Hull, S. I. (1997). Nutritional requirements for growth of uropathogenic Escherichia coli in human urine. Infect. Immun. 65, 1960–1961. doi: 10.1128/iai.65.5.1960-1961.1997

Ipe, D. S., Horton, E., and Ulett, G. C. (2016). The basics of bacteriuria: Strategies of microbes for persistence in urine. Front. Cell. Infect. Microbiol. 6, 14. doi: 10.3389/fcimb.2016.00014

Jacobs, D., Heimbach, D., and Hesse, A. (2001). Chemolysis of struvite stones by acidification of artificial urine: An in vitro study. Scand. J. Urol. Nephrol. 35, 345–349. doi: 10.1080/003655901753224387

Jones, S. M., Yerly, J., Hu, Y., Ceri, H., and Martinuzzi, R. (2007). Structure of Proteus mirabilis biofilms grown in artificial urine and standard laboratory media. FEMS Microbiol. Lett. 268, 16–21. doi: 10.1111/j.1574-6968.2006.00587.x

Juarez, G. E., Mateyca, C., and Galvan, E. M. (2020). Proteus mirabilis outcompetes Klebsiella pneumoniae in artificial urine medium through secretion of ammonia and other volatile compounds. Heliyon 6, e03361. doi: 10.1016/j.heliyon.2020.e03361

Laupland, K. B., Ross, T., Pitout, J. D. D., Church, D. L., and Gregson, D. B. (2007). Community-onset urinary tract infections: A population-based assessment. Infection 35, 150–153. doi: 10.1007/s15010-007-6180-2

Lewis, I. A. (2024). Boundary flux analysis: An emerging strategy for investigating metabolic pathway activity in large cohorts. Curr. Opin. Biotechnol. 85, 103027. doi: 10.1016/j.copbio.2023.103027

Ma, J., Cai, X., Bao, Y., Yao, H., and Li, G. (2018). Uropathogenic Escherichia coli preferentially utilize metabolites in urine for nucleotide biosynthesis through salvage pathways. Int. J. Med. Microbiol. 308, 990–999. doi: 10.1016/j.ijmm.2018.08.006

Norsworthy, A. N., and Pearson, M. M. (2017). From catheter to kidney stone: The uropathogenic lifestyle of Proteus mirabilis. Trends Microbiol. 25, 304–315. doi: 10.1016/j.tim.2016.11.015

Perez, M., Calles-Enríquez, M., Nes, I., Martin, M. C., Fernandez, M., Ladero, V., et al. (2015). Tyramine biosynthesis is transcriptionally induced at low pH and improves the fitness of Enterococcus faecalis in acidic environments. Appl. Microbiol. Biotechnol. 99, 3547–3558. doi: 10.1007/s00253-014-6301-7

Ponce, L. F., Bishop, S. L., Wacker, S., Groves, R. A., and Lewis, I. A. (2024). SCALiR: A web application for automating absolute quantification of mass spectrometry-based metabolomics data. Anal. Chem. 96, 6566–6574. doi: 10.1021/acs.analchem.3c04988

Rasmussen, L. G., Savorani, F., Larsen, T. M., Dragsted, L. O., Astrup, A., and Engelsen, S. B. (2011). Standardization of factors that influence human urine metabolomics. Metabolomics 7, 71–83. doi: 10.1007/s11306-010-0234-7

Roesch, P. L., Redford, P., Batchelet, S., Moritz, R. L., Pellett, S., Haugen, B. J., et al. (2003). Uropathogenic Escherichia coli use D-serine deaminase to modulate infection of the murine urinary tract. Mol. Microbiol. 49, 55–67. doi: 10.1046/j.1365-2958.2003.03543.x

Russo, T. A., Jodush, S. T., Brown, J. J., and Johnson, J. R. (1996). Identification of two previously unrecognized genes (guaA and argC) important for uropathogenesis. Mol. Microbiol. 22, 217–229. doi: 10.1046/j.1365-2958.1996.00096.x

Rydzak, T., Groves, R. A., Zhang, R., Aburashed, R., Pushpker, R., Mapar, M., et al. (2022). Metabolic preference assay for rapid diagnosis of bloodstream infections. Nat. Commun. 13, 2332. doi: 10.1038/s41467-022-30048-6

Sarigul, N., Korkmaz, F., and Kurultak, İ. (2019). A new artificial urine protocol to better imitate human urine. Sci. Rep. 9, 20159. doi: 10.1038/s41598-019-56693-4

Schappert, S. M., and Rechtsteiner, E. A. (2011). Ambulatory medical care utilization estimates for 2007. Vital Health Stat. 13. 13, 1–38.

Segata, N., Haake, S. K., Mannon, P., Lemon, K. P., Waldron, L., Gevers, D., et al. (2012). Composition of the adult digestive tract bacterial microbiome based on seven mouth surfaces, tonsils, throat and stool samples. Genome Biol. 13, R42. doi: 10.1186/gb-2012-13-6-r42

Shaffer, C. L., Zhang, E. W., Dudley, A. G., Dixon, B. R. E. A., Guckes, K. R., Breland, E. J., et al. (2017). Purine biosynthesis metabolically constrains intracellular survival of uropathogenic Escherichia coli. Infect. Immun. 85, e00471–16. doi: 10.1128/IAI.00471-16

Sintsova, A., Smith, S., Subashchandrabose, S., and Mobley, H. L. (2018). Role of ethanolamine utilization genes in host colonization during urinary tract infection. Infect. Immun. 86, e00542–17. doi: 10.1128/IAI.00542-17

Tannock, G. W. (1999). The bowel microflora: An important source of urinary tract pathogens. World J. Urol. 17, 339–344. doi: 10.1007/s003450050158

Thakker, C., Martínez, I., San, K. Y., and Bennett, G. N. (2012). Succinate production in Escherichia coli. Biotechnol. J. 7, 213–224. doi: 10.1002/biot.201100061

Vejborg, R. M., de Evgrafov, M. R., Phan, M. D., Totsika, M., Schembri, M. A., and Hancock, V. (2012). Identification of genes important for growth of asymptomatic bacteriuria Escherichia coli in urine. Infect. Immun. 80, 3179–3188. doi: 10.1128/IAI.00473-12

Virgiliou, C., Theodoridis, G., Wilson, I. D., and Gika, H. G. (2021). Quantification of endogenous amino acids and amino acid derivatives in urine by hydrophilic interaction liquid chromatography tandem mass spectrometry. J. Chromatogr. A 1642, 462005. doi: 10.1016/j.chroma.2021.462005

Wagenlehner, F. M. E., Bjerklund Johansen, T. E., Cai, T., Koves, B., Kranz, J., Pilatz, A., et al. (2020). Epidemiology, definition and treatment of complicated urinary tract infections. Nat. Rev. Urol. 17, 586–600. doi: 10.1038/s41585-020-0362-4

