## Supplementary Material for "Metabolomics survey of uropathogenic bacteria in human urine"

**Supplementary Table 1. List of all bacterial strains used for the metabolomics survey.** The eight most common bacterial species responsible for UTIs were selected and for each species, six strains from diverse origins were chosen for analyses.

| Genus | Species | Strain | Source of Isolation |
|---|---|---|---|
| *Escherichia* | *coli* | K-12 MG1655 | Laboratory control strain |
| *Escherichia* | *coli* | CFT073 | Urine & Blood, Acute pyelonephritis |
| *Escherichia* | *coli* | 1099705 (ATCC BAA-2776) | Urine |
| *Escherichia* | *coli* | NCTC 9001 (ATCC 11775) | Urine, Cystitis |
| *Escherichia* | *coli* | S1-STOOL-BHI-02 (RP14) | Stool |
| *Escherichia* | *coli* | O127:H6 E2348/69 | Stool, Infantile diarrhea |
| *Enterococcus* | *faecalis* | NCTC 775 (ATCC 19433) | Laboratory control strain |
| *Enterococcus* | *faecalis* | Taxo 239 (ATCC 51575) | Laboratory control strain |
| *Enterococcus* | *faecalis* | Portland (ATCC 29212) | Urine |
| *Enterococcus* | *faecalis* | S46-3-BHI-01 (RP3) | Stool |
| *Enterococcus* | *faecalis* | UWH 1921 (ATCC 49532) | Blood |
| *Enterococcus* | *faecalis* | NJ-3 (ATCC 51299) | Peritoneal fluid |
| *Klebsiella* | *quasipneumoniae* (formerly, *pneumoniae*) | K6 (ATCC 700603) | Urine |
| *Klebsiella* | *pneumoniae* | ART2008133 (ATCC BAA-1705) | Urine |
| *Klebsiella* | *pneumoniae* | BK34907 (ATCC BAA-2788) | Urine |
| *Klebsiella* | *pneumoniae* | 17VG-011402-T4-D4 (MG1) | Vagina |
| *Klebsiella* | *pneumoniae* | 931476 (ATCC BAA-2782) | Peritoneal fluid |
| *Klebsiella* | *pneumoniae* | BK34774 (ATCC BAA-2786) | Wound |
| *Pseudomonas* | *aeruginosa* | NCTC 10332 (ATCC 10145) | Laboratory control strain |
| *Pseudomonas* | *aeruginosa* | 1106432 (ATCC BAA-2795) | Urine |
| *Pseudomonas* | *aeruginosa* | Boston 41501 (ATCC 27853) | Blood |
| *Pseudomonas* | *aeruginosa* | 1079232 (ATCC BAA-2794) | Abscess |
| *Pseudomonas* | *aeruginosa* | 1109196 (ATCC BAA-2797) | Endotracheal aspirate |
| *Pseudomonas* | *aeruginosa* | 1124989 (ATCC BAA-2798) | Sputum |
| *Proteus* | *mirabilis* | LRA 08 01 73 (ATCC 35659) | Laboratory control strain |

| | | | |
|---|---|---|---|
| *Proteus* | *mirabilis* | (ATCC 7002) | Urine, Kidney stones |
| *Proteus* | *mirabilis* | CCUG 39506<br>(ATCC BAA-2792) | Urine |
| *Proteus* | *mirabilis* | 927889<br>(ATCC BAA-2791) | Urine |
| *Proteus* | *mirabilis* | NCDC 2059-70<br>(ATCC 25933) | Vagina |
| *Proteus* | *mirabilis* | CCHMC1<br>(ATCC BAA-856) | Clinical isolate |
| *Streptococcus* | *agalactiae* | 330434371 | Clinical isolate |
| *Streptococcus* | *agalactiae* | 330438073 | Clinical isolate |
| *Streptococcus* | *agalactiae* | 330438068 | Clinical isolate |
| *Streptococcus* | *agalactiae* | 330435372 | Clinical isolate |
| *Streptococcus* | *agalactiae* | 330438078 | Clinical isolate |
| *Streptococcus* | *agalactiae* | 330438048 | Clinical isolate |
| *Staphylococcus* | *aureus* | FDA<br>(ATCC 29737) | Laboratory control strain |
| *Staphylococcus* | *aureus* | Seattle 1945<br>(ATCC 25923) | Clinical isolate |
| *Staphylococcus* | *aureus* | FDA 209<br>(ATCC 6538) | Lesion |
| *Staphylococcus* | *aureus* | UT 32<br>(ATCC BAA-977) | Wound |
| *Staphylococcus* | *aureus* | Wichita<br>(ATCC 29213) | Wound |
| *Staphylococcus* | *aureus* | UT 25<br>(ATCC BAA-976) | Tracheal aspirate |
| *Staphylococcus* | *saprophyticus* | Vitek #8935<br>(ATCC BAA-750) | Laboratory control strain |
| *Staphylococcus* | *saprophyticus* | API 222<br>(ATCC 49453) | Laboratory control strain |
| *Staphylococcus* | *saprophyticus* | AmMS 143<br>(ATCC 49907) | Laboratory control strain |
| *Staphylococcus* | *saprophyticus* | API 1101<br>(ATCC 35552) | Laboratory control strain |
| *Staphylococcus* | *saprophyticus* | LRA 27.02.80<br>(ATCC 43867) | Laboratory control strain |
| *Staphylococcus* | *saprophyticus* | NCTC 7292<br>(ATCC 15305) | Urine |

**Supplementary Table 2. Concentrations of common metabolites in pooled human urine that was used as growth medium for *in vitro* bacterial cultures.** Common metabolites (amino acids, amino acid derivatives, nucleic acids, carbohydrates, vitamins) were quantified by running three replicates of the urine stock alongside a series of mixed standards. Central carbon metabolites were grouped as "carbohydrates". Key consumed metabolites (>1 μM by at least one species) were marked with asterisk (*) and these metabolites should be added to artificial urine to better mimic human urine.

| Metabolite | Average Concentration (μM) (± standard deviation) |
|---|---|
| **Amino Acids** | |
| Arginine* | 20.68 ± 0.62 |
| Asparagine* | 96.59 ± 1.45 |
| Aspartate | 53.50 ± 25.77 |
| Cysteine* | 192.70 ± 22.10 |
| Glutamic Acid* | 29.90 ± 0.78 |
| Glutamine* | 344.15 ± 7.72 |
| Glycine | 681.11 ± 84.62 |
| Histidine | 476.59 ± 20.60 |
| Isoleucine* | 44.85 ± 9.59 |
| Leucine* | 90.41 ± 10.84 |
| Lysine* | 69.83 ± 3.22 |
| Methionine* | 13.81 ± 2.79 |
| Phenylalanine* | 67.70 ± 4.05 |
| Proline* | 51.11 ± 15.19 |
| Serine* | 230.65 ± 3.53 |
| Threonine* | 126.15 ± 2.87 |
| Tryptophan* | 24.22 ± 1.19 |
| Tyrosine* | 50.60 ± 3.84 |
| Valine* | 68.96 ± 6.76 |
| **Amino Acid Derivatives** | |
| N-Acetyl-Aspartic Acid* | 58.25 ± 1.21 |
| N-Acetyl-Glutamic Acid* | 22.29 ± 0.22 |
| N-Acetyl-Glutamine | 13.46 ± 0.80 |
| N-Acetyl-Glycine | 4.81 ± 0.32 |
| N-Acetyl-Leucine | 7.15 ± 0.18 |
| N-Acetyl-Methionine | 0.03 ± 0.02 |
| N-Acetyl-Phenylalanine | 1.11 ± 0.05 |
| Acetyl-Threonine | 48.26 ± 1.05 |
| Agmatine | 0.01 ± 0.01 |
| Carnosine | 22.03 ± 0.76 |
| Choline | 19.06 ± 3.47 |
| Citrulline* | 6.19 ± 1.16 |
| Creatinine | 8613.35 ± 187.71 |
| Cysteic Acid | 0.00 ± 0.00 |
| Cystine* | 28.56 ± 1.72 |
| Hippurate* | 1320.70 ± 26.10 |

| | |
|---|---|
| Homocysteine | 0.02 ± 0.04 |
| 4-Hydroxyproline | 3.03 ± 1.21 |
| O-Phospho-Serine | 20.66 ± 11.81 |
| Ornithine* | 10.06 ± 1.68 |
| Methionine Sulfoxide* | 7.47 ± 1.41 |
| Shikimate | 0.68 ± 0.03 |
| Spermine | 45.07 ± 20.4 |
| Taurine* | 345.67 ± 5.63 |
| Tyramine | 0.82 ± 0.10 |
| Urocanate | 2.31 ± 0.11 |
| **Nucleic Acids** | |
| Adenine | 0.15 ± 0.02 |
| Adenosine | 1.72 ± 0.03 |
| Adenosine Monophosphate | 0.00 ± 0.00 |
| Allantoin | 72.83 ± 1.74 |
| Cytidine | 0.77 ± 0.05 |
| Deoxyadenosine | 0.03 ± 0.00 |
| Deoxycytidine | 0.05 ± 0.09 |
| Deoxyguanosine* | 12.09 ± 0.72 |
| Deoxyuridine | 0.02 ± 0.01 |
| Dihydroorotate | 0.44 ± 0.02 |
| Guanine | 0.84 ± 0.03 |
| Guanosine | 0.38 ± 0.02 |
| Hypoxanthine* | 48.98 ± 1.50 |
| Inosine* | 1.66 ± 0.02 |
| 3-Methyladenine | 0.06 ± 0.01 |
| Orotate | 7.69 ± 0.34 |
| Thymidine | 0.04 ± 0.04 |
| Thymine | 0.69 ± 0.04 |
| Uracil | 0.01 ± 0.01 |
| Uridine | 0.69 ± 0.01 |
| Xanthine* | 10.67 ± 0.25 |
| Xanthosine* | 4.85 ± 0.18 |
| **Carbohydrates** | |
| Acetoacetate | 3.59 ± 0.74 |
| N-Acetyl-Glucosamine* | 64.00 ± 2.46 |
| Aconitate* | 185.98 ± 2.56 |
| Arabitol* | 221.31 ± 2.03 |
| Fructose 1,6-Biphosphate | 35.57 ± 14.39 |
| Fumarate* | 22.92 ± 1.08 |
| Glucose* | 248.60 ± 3.18 |
| Glucose 6-Phosphate | 7.06 ± 2.36 |
| Glycerol 3-Phosphate* | 26.89 ± 3.42 |
| Itaconate* | 1.54 ± 0.03 |
| Mevalonate | 0.08 ± 0.01 |

| | |
|---|---|
| Ribose 5-Phosphate | 7.55 ± 3.90 |
| Succinate* | 328.63 ± 3.71 |
| Trehalose | 1.21 ± 0.05 |
| **Vitamins** | |
| Biotin | 0.07 ± 0.01 |
| Nicotinamide | 0.42 ± 0.01 |
| Nicotinate | 0.04 ± 0.01 |
| Pantothenic Acid | 11.65 ± 0.09 |
| Pyridoxine | 1.14 ± 0.56 |
| Riboflavin | 0.09 ± 0.02 |
| Thiamine | 0.96 ± 0.34 |

**Supplementary Table 3. Metabolite concentration changes in bacterial urine cultures.** Average concentration changes of various common metabolites after *in vitro* four-hour growth course in human urine, subtracted against urine controls. Metabolite concentration increases (in red) and decreases (in blue) based on two-sample t-tests ($p<0.05$) are stated in micromolar (μM). Dash (-) indicates there was no difference ($p>0.05$, two-sample t-test) between the start and end of the growth course.

| Metabolites | *E. coli* (n=6) | *K. pneumoniae* (n=6) | *P. mirabilis* (n=6) | *P. aeruginosa* (n=6) | *E. faecalis* (n=6) | *S. agalactiae* (n=6) | *S. aureus* (n=6) | *S. saprophyticus* (n=6) |
|---|---|---|---|---|---|---|---|---|
| **Amino Acids** | | | | | | | | |
| Arginine | -5.76 | -5.11 | -15.03 | -15.10 | -6.70 | -6.24 | -6.98 | -5.15 |
| Asparagine | -67.53 | -36.94 | -63.15 | -63.34 | - | - | - | -6.38 |
| Aspartate | - | - | - | - | - | - | - | - |
| Cysteine | -37.18 | - | -72.09 | - | - | - | - | -26.57 |
| Glutamic Acid | -6.39 | - | - | -7.50 | +88.99 | +52.25 | - | +47.35 |
| Glutamine | -148.74 | -129.18 | -91.73 | -281.57 | - | - | -88.07 | -83.67 |
| Glycine | - | - | - | - | - | +365.40 | +622.95 | - |
| Histidine | - | - | +97.38 | - | - | +35.40 | - | +22.53 |
| Isoleucine | -15.88 | -15.16 | -21.05 | -15.51 | - | - | -19.58 | -17.81 |
| Leucine | -29.19 | - | -39.10 | - | - | - | -25.24 | - |
| Lysine | - | -42.55 | - | - | - | +76.40 | - | - |
| Methionine | -2.30 | -2.31 | -2.24 | - | - | - | -2.11 | -1.27 |
| Phenylalanine | -10.97 | -7.66 | -35.10 | - | - | +27.37 | - | - |
| Proline | - | -19.98 | - | -16.24 | - | - | - | -20.01 |
| Serine | -172.12 | -187.79 | -182.30 | -48.86 | - | - | -57.06 | -77.71 |
| Threonine | -77.38 | -38.13 | -68.15 | - | - | - | -70.29 | -49.58 |
| Tryptophan | -19.44 | - | -12.39 | - | - | - | -1.59 | -2.71 |
| Tyrosine | -10.89 | -4.63 | -24.66 | - | -15.35 | - | -14.37 | -11.66 |
| Valine | -39.66 | -36.56 | -37.10 | -32.27 | - | - | -34.20 | -24.16 |
| **Amino Acid Derivatives** | | | | | | | | |
| N-Acetyl-Aspartic Acid | - | - | -49.40 | - | - | - | -3.08 | -5.71 |
| N-Acetyl-Glutamic Acid | +2.89 | - | +2.74 | - | - | - | - | -1.88 |
| N-Acetyl-Glutamine | - | - | - | - | - | - | -0.82 | - |
| N-Acetyl-Glycine | +0.76 | - | - | - | - | - | +0.63 | -0.38 |
| N-Acetyl-Leucine | - | - | - | - | - | - | -0.35 | - |
| N-Acetyl-Methionine | - | - | - | - | - | - | - | - |
| N-Acetyl-Phenylalanine | - | - | - | - | - | - | -0.07 | - |
| Acetyl-Threonine | - | - | - | - | - | - | - | - |
| Agmatine | +0.04 | +0.02 | - | - | - | - | - | - |

| | | | | | | | | |
|---|---|---|---|---|---|---|---|---|
| Carnosine | - | - | +2.86 | - | +1.25 | +1.79 | - | - |
| Choline | - | - | +7.53 | - | - | - | - | - |
| Citrulline | -2.54 | -2.38 | +1.25 | +3.33 | - | - | - | - |
| Cysteic Acid | - | - | - | - | - | - | - | - |
| Cystine | -7.68 | -4.53 | -17.80 | - | - | - | - | - |
| Hippurate | - | - | +166.59 | - | - | - | -1363.05 | -203.95 |
| Homocysteine | - | - | - | - | - | - | - | - |
| 4-Hydroxyproline | +6.64 | +7.99 | +10.06 | - | - | - | - | - |
| O-Phospho-Serine | - | - | - | - | - | - | - | - |
| Ornithine | -2.54 | - | - | - | +13.61 | - | +2.65 | - |
| Methionine Sulfoxide | -2.14 | -1.93 | -2.84 | - | - | - | -1.10 | - |
| Shikimate | - | - | - | - | - | - | - | - |
| Spermine | -0.27 | - | +0.52 | -0.50 | - | -0.57 | -0.89 | -0.95 |
| Taurine | - | +23.51 | +37.18 | - | - | - | -21.54 | -20.84 |
| Tyramine | - | - | - | - | +2.50 | - | - | - |
| Urocanate | - | +1.25 | +0.18 | - | - | - | - | - |
| **Nucleic Acids** | | | | | | | | |
| Adenine | -0.08 | - | +1.80 | +0.14 | - | +1.60 | +1.76 | +0.70 |
| Adenosine | -0.88 | -0.85 | -0.67 | - | -0.85 | -0.88 | -0.87 | -0.91 |
| Adenosine Monophosphate | - | - | - | - | - | - | - | - |
| Allantoin | +13.61 | +12.63 | +19.97 | +9.18 | - | - | +7.42 | - |
| Cytidine | -0.62 | -0.60 | -0.55 | - | -0.58 | - | -0.57 | -0.27 |
| Deoxyadenosine | -0.02 | -0.02 | -0.01 | - | -0.02 | -0.02 | -0.02 | -0.02 |
| Deoxycytidine | - | - | - | - | - | - | - | - |
| Deoxyguanosine | -10.86 | -8.78 | -7.70 | - | -10.37 | -8.68 | -7.71 | -3.70 |
| Deoxyuridine | - | - | - | - | - | - | - | - |
| Dihydroorotate | +0.41 | +0.82 | -0.12 | +0.46 | - | -0.08 | - | - |
| Guanine | - | +1.01 | +1.45 | -0.67 | +0.66 | +0.43 | - | +0.26 |
| Guanosine | -0.37 | -0.34 | -0.23 | - | - | -0.17 | -0.20 | -0.16 |
| Hypoxanthine | -18.57 | -12.28 | -10.62 | -29.74 | - | - | -11.09 | -6.72 |
| Inosine | -0.99 | -0.86 | -1.12 | - | -0.82 | -0.86 | -0.73 | -0.66 |
| 3-Methyladenine | +0.15 | +0.16 | +0.20 | +0.06 | - | +0.22 | +0.20 | +0.32 |

| | | | | | | | | |
|---|---|---|---|---|---|---|---|---|
| Orotate | +0.67 | - | +2.65 | - | - | - | - | - |
| Thymidine | - | -0.06 | - | - | -0.07 | -0.05 | -0.09 | -0.09 |
| Thymine | - | - | +203.62 | +0.45 | - | - | - | - |
| Uracil | - | - | - | - | - | - | - | - |
| Uridine | -0.75 | -0.74 | -0.70 | - | -0.75 | -0.75 | -0.68 | -0.70 |
| Xanthine | -3.60 | -7.10 | -3.65 | - | - | - | -3.91 | -1.53 |
| Xanthosine | - | - | +0.48 | - | - | -1.71 | - | -0.36 |
| **Carbohydrates** | | | | | | | | |
| Acetoacetate | - | - | - | - | - | - | - | - |
| N-Acetyl-Glucosamine | -47.55 | -40.74 | -26.85 | - | -49.46 | -43.89 | -45.28 | -15.75 |
| Aconitate | - | - | - | -35.14 | - | - | -13.30 | -16.88 |
| Arabitol | +21.45 | -95.36 | +33.64 | - | - | - | - | -16.34 |
| Fructose 1,6-Bisphosphate | - | - | - | - | - | - | - | - |
| Fumarate | - | - | -21.23 | -22.01 | - | - | - | - |
| Glucose | -221.88 | -224.92 | -211.50 | - | -212.18 | -212.05 | -207.17 | -196.37 |
| Glucose 6-Phosphate | - | +1.93 | - | - | - | - | - | - |
| Glycerol 3-Phosphate | -17.08 | -16.74 | - | - | - | - | -10.58 | -1.61 |
| Itaconate | - | - | - | -0.76 | -1.08 | - | - | - |
| Mevalonate | +0.11 | - | +0.03 | - | - | - | - | +0.04 |
| Ribose 5-Phosphate | +1.91 | +1.54 | +1.81 | - | - | +1.37 | - | - |
| Succinate | +1003.44 | - | -300.13 | -346.06 | - | - | +184.60 | +47.74 |
| Trehalose | -0.44 | -0.56 | - | - | - | - | -0.41 | -0.26 |
| **Vitamins** | | | | | | | | |
| Biotin | - | - | - | - | - | - | - | - |
| Nicotinamide | -0.37 | -0.38 | -0.37 | -0.37 | -0.37 | -0.30 | -0.29 | -0.35 |
| Nicotinate | +0.43 | +0.43 | +0.71 | +0.38 | +0.15 | +0.35 | - | +0.34 |
| Pantothenic Acid | - | - | - | - | - | - | -0.50 | - |
| Pyridoxine | - | - | - | - | +0.06 | +0.02 | - | - |
| Riboflavin | +0.06 | - | -0.03 | - | - | - | - | -0.03 |
| Thiamine | - | - | -0.06 | - | - | - | -0.02 | - |

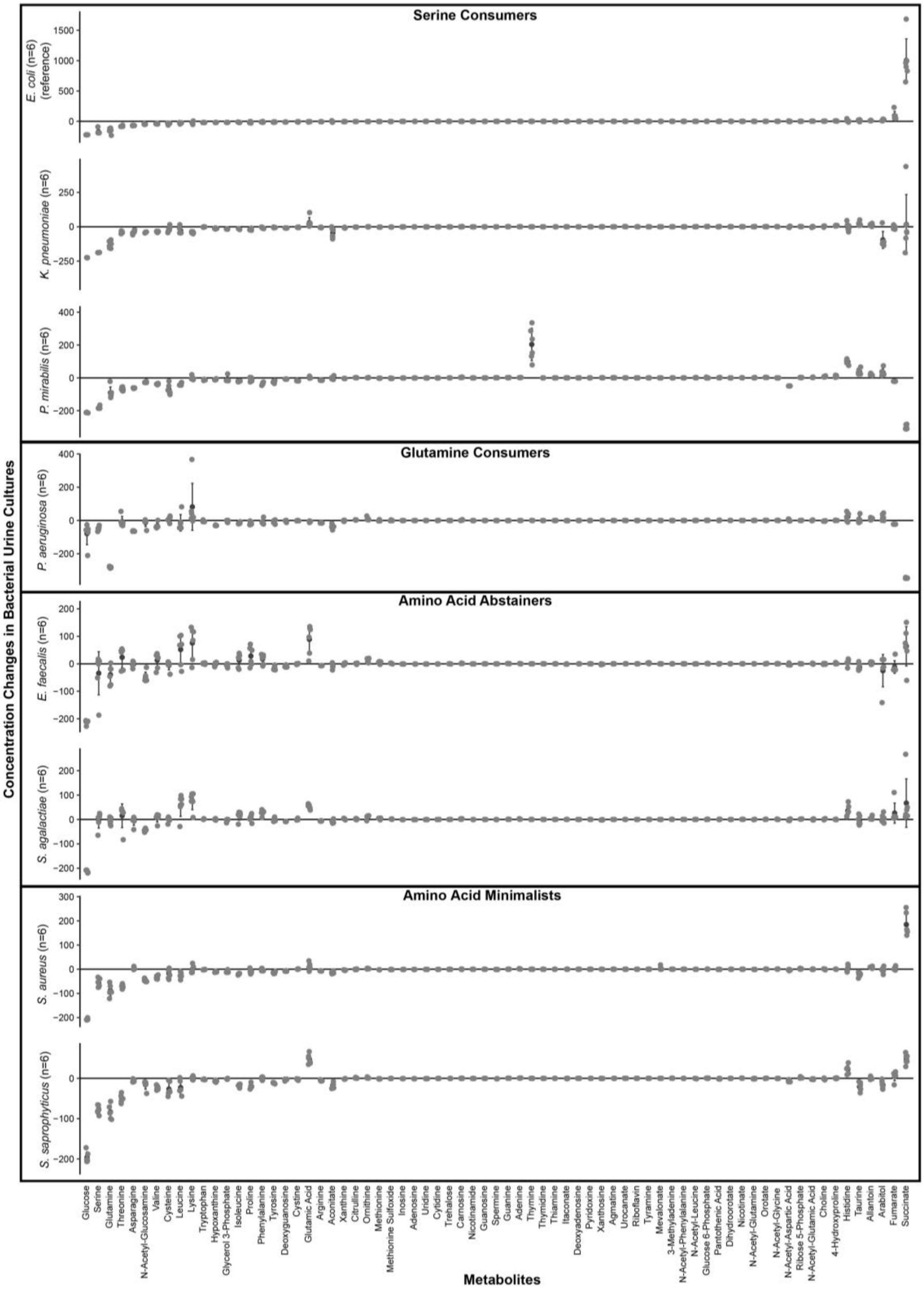


**Supplementary Figure 1. Metabolite concentration changes in *in vitro* urine cultures of the eight most common uropathogenic species sorted by metabolic clade.** Metabolites with concentration changes were sorted by most consumed to most produced in *E. coli* cultures as a reference for all other species. Error bars represent one standard deviation.

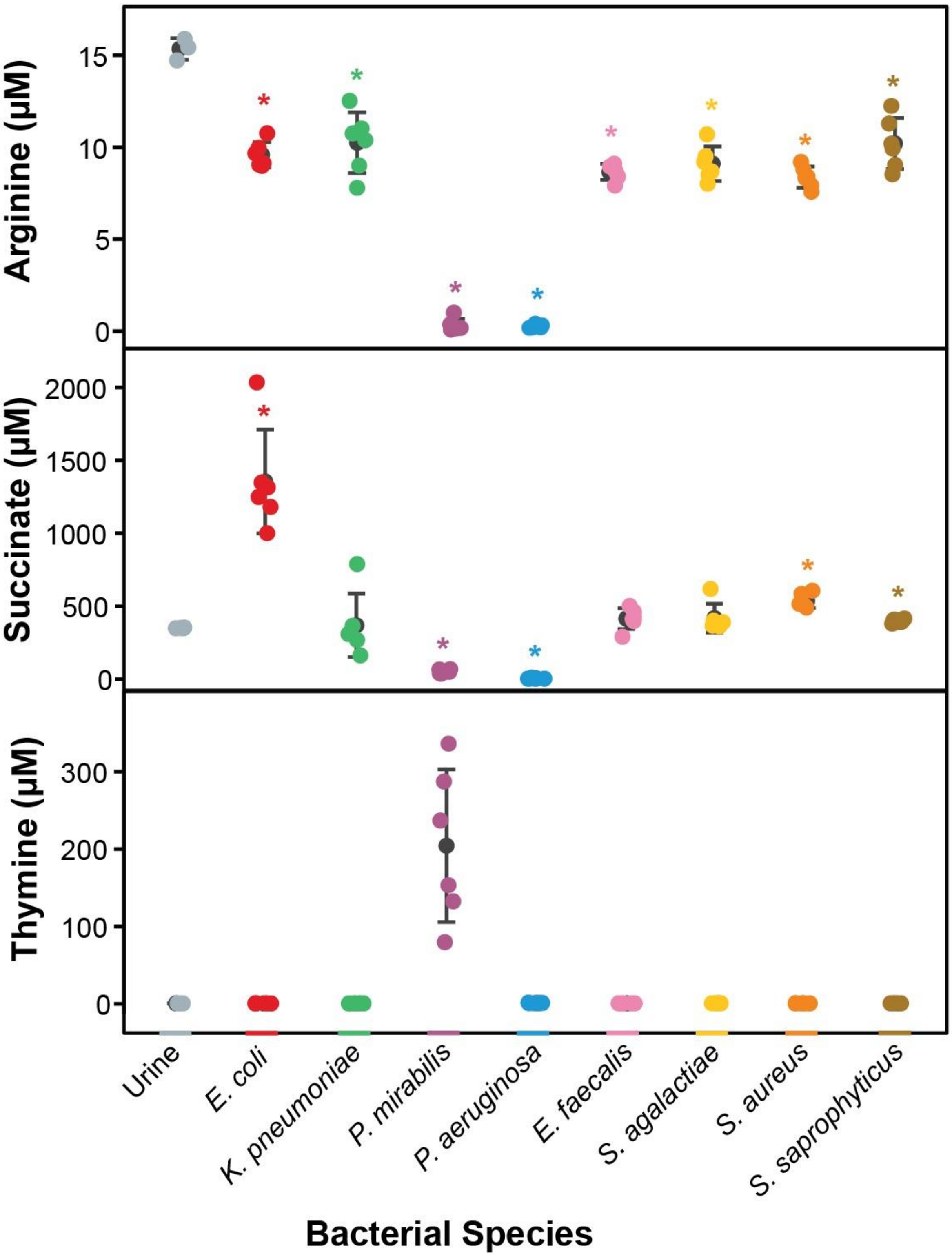


**Supplementary Figure 2. Select metabolic phenotypes observed in specific species during *in vitro* growth in urine.** Concentrations of arginine, succinate and thymine in urine cultures of eight bacterial species (six strains per species). Statistical significance is denoted by an asterisk (*, two-sample t-test with Bonferroni correction $\alpha$=0.00625). Error bars represent one standard deviation.

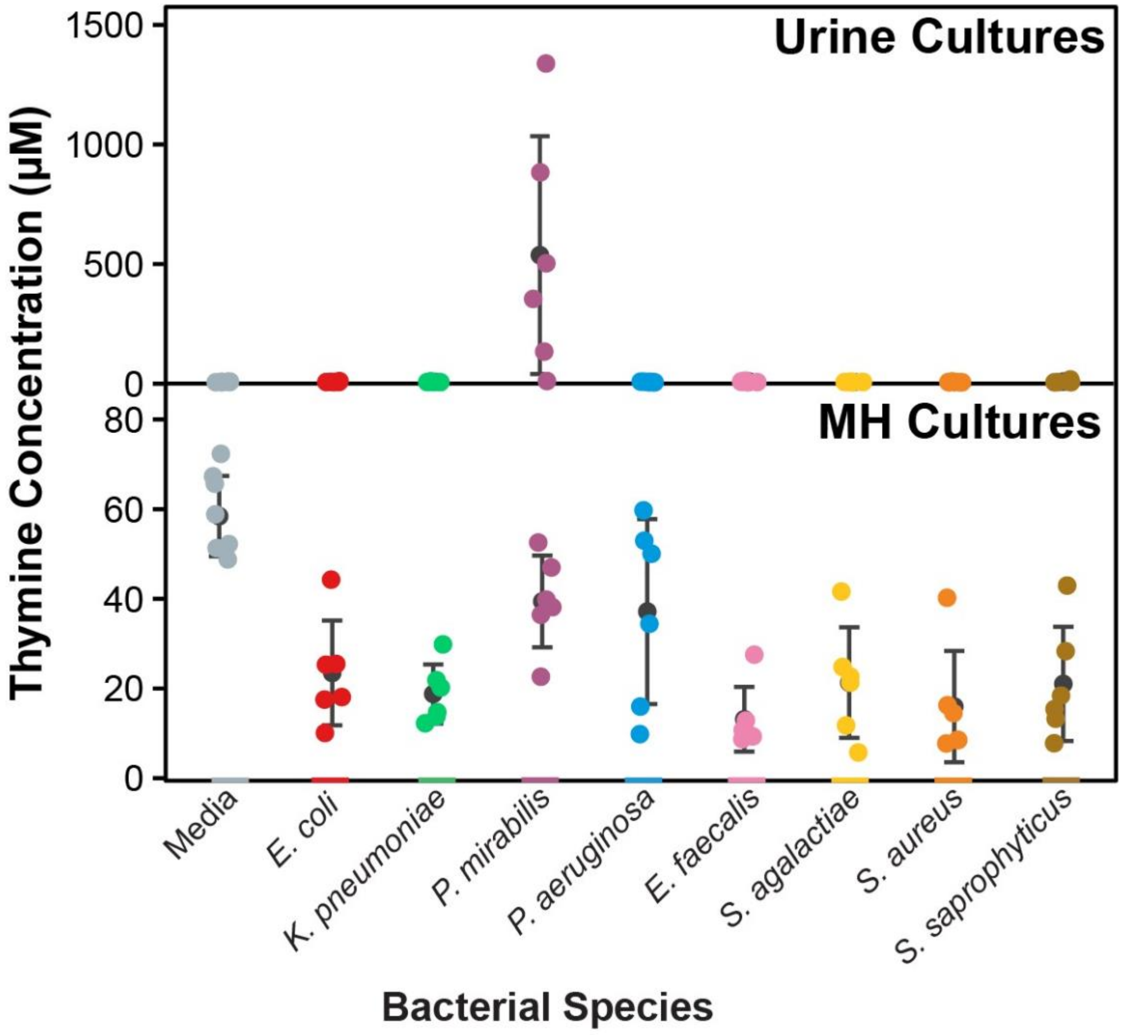


**Supplementary Figure 3. Thymine concentrations in *in vitro* urine cultures compared to Mueller-Hinton (MH) cultures.** Concentrations of thymine in urine and MH cultures of eight bacterial species (six strains per species). Error bars represent one standard deviation.

**Supplementary Table 4. Chi-square analysis of the observed species prevalence in a cohort of polymicrobial UTIs**. Based on a cohort of patient urine samples sent for clinical urine culture over the course of a week, 5.5% (81/1462) of growth-positive urine samples contained two causative species. The top three most prevalence species pairings each underwent a chi-square goodness-of-fit test to determine if the pairing occurred by chance or not. The expected occurrences were obtained through probability analysis of all species in the polymicrobial UTI cohort. df, degrees of freedom.

| Species 1 | Individual Prevalence (%) | Species 2 | Individual Prevalence (%) | Expected Combined Occurrence (out of 81) | Observed Combined Occurrence (out of 81) | Chi-Square Statistic | P-Value (df=1) |
|---|---|---|---|---|---|---|---|
| *Escherichia coli* | 34.0 | *Enterococcus faecalis* | 22.8 | 15.4 | 26 | 9.1 | 0.0026 |
| *Escherichia coli* | 34.0 | *Klebsiella pneumoniae* | 8.0 | 5.4 | 7 | 0.5 | 0.4756 |
| *Escherichia coli* | 34.0 | *Streptococcus viridans* | 4.3 | 2.9 | 7 | 6.0 | 0.0145 |
| *Escherichia coli* | 34.0 | *Aerococcus urinae* | 3.1 | 2.1 | 4 | - | - |
| *Klebsiella pneumoniae* | 8.0 | *Enterococcus faecalis* | 22.8 | 3.6 | 4 | - | - |
| *Enterococcus faecalis* | 22.8 | *Morganella morganii* | 1.9 | 0.8 | 2 | - | - |
| *Escherichia coli* | 34.0 | *Staphylococcus sp.* (coagulase-negative) | 1.2 | 0.8 | 2 | - | - |
| *Escherichia coli* | 34.0 | *Enterobacter cloacae* | 2.5 | 1.7 | 2 | - | - |
| *Escherichia coli* | 34.0 | *Enterococcus faecium* | 3.7 | 2.5 | 2 | - | - |
| *Klebsiella pneumoniae* | 8.0 | *Citrobacter freundii* | 3.1 | 0.5 | 2 | - | - |
| *Pseudomonas aeruginosa* | 3.1 | *Enterococcus faecalis* | 22.8 | 1.4 | 2 | - | - |
| *Aerococcus urinae* | 3.1 | *Aerococcus sanguinicola* | 1.9 | 0.1 | 1 | - | - |
| *Enterobacter cloacae* | 2.5 | *Citrobacter koseri* | 0.6 | 0.0 | 1 | - | - |
| *Enterococcus faecalis* | 22.8 | *Citrobacter freundii* | 3.1 | 1.4 | 1 | - | - |
| *Enterococcus faecalis* | 22.8 | *Lactobacillus sp.* | 0.6 | 0.3 | 1 | - | - |
| *Enterococcus faecium* | 3.7 | *Candida albicans* | 0.6 | 0.0 | 1 | - | - |
| *Enterococcus faecium* | 3.7 | *Citrobacter freundii* | 3.1 | 0.2 | 1 | - | - |
| *Enterococcus faecium* | 3.7 | *Hafnia alvei* | 0.6 | 0.0 | 1 | - | - |
| *Escherichia coli* | 34.0 | *Aerococcus viridans* | 0.6 | 0.4 | 1 | - | - |
| *Escherichia coli* | 34.0 | *Citrobacter amalonaticus* | 0.6 | 0.4 | 1 | - | - |
| *Escherichia coli* | 34.0 | *Enterococcus raffinosus* | 0.6 | 0.4 | 1 | - | - |
| *Escherichia coli* | 34.0 | *Klebsiella oxytoca* | 1.9 | 1.2 | 1 | - | - |

| | | | | | | | |
|---|---|---|---|---|---|---|---|
| *Escherichia coli* | 34.0 | *Streptococcus agalactiae* | 0.6 | 0.4 | 1 | - | - |
| *Klebsiella oxytoca* | 1.9 | *Morganella morganii* | 1.9 | 0.1 | 1 | - | - |
| *Proteus mirabilis* | 1.2 | *Aerococcus sanguinicola* | 1.9 | 0.0 | 1 | - | - |
| *Proteus mirabilis* | 1.2 | *Citrobacter freundii* | 3.1 | 0.1 | 1 | - | - |
| *Providencia rettgeri* | 0.6 | *Aerococcus sanguinicola* | 1.9 | 0.0 | 1 | - | - |
| *Pseudomonas aeruginosa* | 3.1 | *Enterobacter cloacae* | 2.5 | 0.2 | 1 | - | - |
| *Pseudomonas aeruginosa* | 3.1 | *Enterococcus faecium* | 3.7 | 0.2 | 1 | - | - |
| *Pseudomonas aeruginosa* | 3.1 | *Proteus vulgaris* | 0.6 | 0.0 | 1 | - | - |
| *Staphylococcus aureus* | 1.2 | *Enterococcus faecalis* | 22.8 | 0.6 | 1 | - | - |
| *Staphylococcus aureus* | 1.2 | *Klebsiella oxytoca* | 1.9 | 0.0 | 1 | - | - |
